# Genomic Biomarker Heterogeneities Between SARS-CoV-2 and COVID-19

**DOI:** 10.1101/2022.01.13.476223

**Authors:** Zhengjun Zhang

## Abstract

Genes functionally associated with SARS-CoV-2 infection and genes functionally related to COVID-19 disease can be different, whose distinction will become the first essential step for successfully fighting against the COVID-19 pandemic. Unfortunately, this first step has not been completed in all biological and medical research. Using a newly developed max-competing logistic classifier, two genes, ATP6V1B2 and IFI27, stand out to be critical in transcriptional response to SARS-CoV-2 infection with differential expressions derived from NP/OP swab PCR. This finding is evidenced by combining these two genes with one another gene in predicting disease status to achieve better-indicating accuracy than existing classifiers with the same number of genes. In addition, combining these two genes with three other genes to form a five-gene classifier outperforms existing classifiers with ten or more genes. These two genes can be critical in fighting against the COVID-19 pandemic as a new focus and direction with their exceptional predicting accuracy. Comparing the functional effects of these genes with a five-gene classifier with 100% accuracy identified and tested from blood samples in our earlier work, genes and their transcriptional response and functional effects to SARS-CoV-2 infection and genes and their functional signature patterns to COVID-19 antibody are significantly different, which can be interpreted as the former is the point of a phenomenon, and the latter is the essence of the disease. We will use a total of fourteen cohort studies (including breakthrough infections and omicron variants) with 1481 samples to justify our results. Such significant findings can help explore the causal and pathological clue between SARS-CoV-2 infection and COVID-19 disease and fight against the disease with more targeted genes, vaccines, antiviral drugs, and therapies.

## 1 Introduction

The fluctuations in infection rates of the COVID-19 pandemic have been like sea waves, with many small ones and several big ones in the past two years. In the meantime, variants of SARS-CoV-2 have emerged and put scientists and medical practitioners on high alert all the time, and many problems have remained unanswered^1;2;3;4;5;6;7;8;9;10;11^. In addition, there have been new concerns with COVID-19 disease, e.g., SARS-CoV-2 enters the brain^12^, COVID-19 vaccines complicate mammograms^13^, memory loss, and ‘brain fog’^14^, amongst others. However, these new concerns are observational and experimental laboratory outcomes, and the genetic bases of those phenomena have not been properly assessed due to a lack of adequate analytical methods to link COVID-19 to the situations. Regarding samples assessed by gene expression profiling, the literature didn’t point out the significant difference between samples with differential expressions derived from nasopharyngeal (NP) and oropharyngeal (OP) swabs PCR and samples derived from whole blood, as the majority of research work focused on individual genes’ expression levels, especially high expression values. Zhang^15^ first applied an innovative algorithm to analyze 126 whole blood samples from COVID-19-positive and COVID-19-negative patients^16^ and reported five critical genes and their competing classifiers, leading to 100% accuracy in classifying all hospitalized patients, including ICU patients, to their respective groups. Zhang^17^ further develops a mathematical and biological equivalence between COVID-19 and five critical genes and proves the existence of at least three transcriptomic data signature patterns and at least seven subtypes. This paper studies gene expression data drawn from NP/OP swab PCR-tested samples with COVID-19 positives and negatives. Surprisingly, we find that the functional effects of those five critical genes, ABCB6, KIAA1614, MND1, SMG1, RIPK3, found in Zhang^15;17^, no longer play a decisive role in NP/OP swap PCR samples. At first glance, this observation seems not useful at all, or it even brings doubts about the study methodology, genomics, and epigenetics. A careful thought confirms that this observation perfectly suggests the relationship between whole blood samples and NP/OP swap PCR samples. The former (whole blood) stands for the essence of the disease, while the latter (NP/OP) stands for the point of the phenomenon. Metaphorically, let’s consider water quality and mineral examination with samples from the deep sea and samples from the shoreside. The samples from the deep sea represent the meta contents and functions of the sea, while the samples from the shoreside contain likely polluted contents from the bank. Also, the structures of the deep sea have changed along sea waves. As a result, samples from the deep sea and samples from the shoreside will provide very different information, with an exception that the whole sea is evenly cleaned or polluted. Here, deep-sea samples correspond to whole blood samples, while shoreside samples correspond to NP/OP swaps PCR samples, which explains the significant difference inferred from the study in Zhang^15;17^and this study.

On the other hand, our new finding calls forth an old question: treat the symptoms, cure the root cause, or both. Zhang^17^ argues that the existence of a genomic signature pattern has to be solved to end the disease, i.e., it is about to cure the root cause. On the other hand, this paper is about treating the symptoms. These two types of research reinforce each other, and both are important to current studies of diseases (any types).

The studies^15;17;18;19;20;21^ applied an innovative algorithm to study classifications of COVID-19 patients, breast cancer patients, lung cancer patients, colorectal cancer patients, and liver cancer patients and gained the highest accuracy (nearly 100%) among eleven different study cohorts with thousands of patients. The high accuracy establishes a mathematical and biological equivalence between the formed classifiers and the disease, which shows that the study method is effective, informative, and robust. These applications are advanced as they lead to new, interpretable, and insightful functional effects of genes linked to the diseases. Using the breast cancer study^18^ as an example, it was found that the known eight famous genes -- BRCA1, BRCA2, PALB2, BARD1, RAD51C, RAD51D, ATM – in breast cancer research and practice are actually leading to very low accuracy in predicting breast cancer status at the stage of diagnosis. Table 6 in the paper^18^ demonstrated that any of these eight genes are very weakly correlated, at most 0.341, with the high-performance biomarkers/genes identified in the study^18^. The findings using our new innovative approach (max logistic classifier) can be the key factor in achieving breakthroughs against the diseases. Due to the limitation of the existing analysis methods and the limited knowledge of the diseases, the fundamental functional effects of genes associated with the disease couldn’t be quested even the truth in the collected data has existed for a long time, and the chances of discovering the truth have been wasted. Conducting new experiments, producing new data, and applying the same analysis methods just like repeating, making the same errors of finding suboptimal (even sometimes misleading) answers. For example, it has been reported that the flu vaccine was only about 16 percent effective, the C.D.C. says. Though the efficacy of a vaccine involves many factors, e.g., the rate of virus mutation, recombination, or aspects of its biological cycle, other than by technical aspects of classification or design studies, identifying fundamental genomic/genetic gene-gene interactions can be intrinsic. This paper uses the innovative method to study differential expressions of human upper respiratory tract gene expressions from 93 COVID-19 positive patients and 141 patients having other acute respiratory illnesses with or without viral infections^22^, and to study host gene expression among RNA-sequencing profiles of nasopharyngeal swabs from 430 individuals with SARS-CoV-2 infection and 54 negative controls^23^. In addition to these two datasets, we will study additional twelve datasets, including blood-sampled datasets and Omicron variants. Details, including how to do crossvalidation with heterogeneous datasets that haven’t been studied, will be discussed in Section 3. Using the first dataset, we identify two genes, ATP6V1B2 and IFI27, critical in transcriptional response to SARS-CoV-2 infection. The gene IFI27 was also identified in Mick et al. (2020)^22^ but did not enter their final classifiers. In the analysis of the first dataset, a combination of these two genes with RIPK3^15^ can lead to an overall accuracy of 87.2%, a sensitivity of 76.3%, and a specificity of 94.3%, and a combination of these two genes with one of these three genes, BTN3A1, SERTAD4, EPSTI1, can lead to an overall accuracy of 89.74%, the sensitivities ranged between 89.25~93.55%, and the specificities between 87.24~90.12%, which are higher than the classifiers in the literature. Using these two genes and one other gene together can easily get overall accuracy between 87.2% and 89.74%, revealing that these two genes can be fundamental. Combining all these five genes can get to an overall accuracy of 91.88%, a sensitivity of 94.62%, and a specificity of 90.08%, higher than the classifiers with 10 genes or more in the literature. Many other combinations will be illustrated in Data Section. These performance results from different combinations indicate that COVID-19 can have many different variants. Unlike the studies in Zhang^15;17^, the accuracy from any combinations applied to NP/OP swaps PCR gene expressions hasn’t been up to 100%. There are three possible reasons, e.g., 1) samples themselves can be false positive or false negative from NP/OP swaps PCR tests; 2) sample signals were weak, and counts were inaccurate; 3) experimental conditions vary. Nevertheless, given the superior performance in the first dataset, the findings shed light on studying SARS-CoV-2 and infections.

These two critical genes ATP6V1B2 (ATPase H+ Transporting V1 Subunit B2) and IFI27 (Interferon Alpha Inducible Protein 27) had previously been reported to be associated with several diseases. For example, de novo mutation in ATP6V1B2 was found to impair lysosome acidification and cause dominant deafness-onychodystrophy syndrome^24^, while IFI27 was found to discriminate between influenza and bacteria in patients with suspected respiratory infection^24^, among others. In addition, a most recent study found that SARS-CoV-2 appeared to persist in organs throughout the body for months^26^.

The significant differences in gene functional effects, gene-gene interactions, and gene-variants interactions between whole blood sampled gene expressions and NP/OP swap PCR sampled gene expressions reveal that ATP6V1B2 and IFI27 are associated with SARS-CoV-2, which points to a new optimal direction of developing more effective vaccines and antiviral drugs. On the other hand, the functional effects of ABCB6, KIAA1614, MND1, SMG1, RIPK3 can be critical to understanding the disease.

The contribution of this paper includes: 1) signifying the genomic difference between NP/OP swap PCR samples and whole blood samples (hospitalized patients); 2) identifying single-digit critical genes (ATP6V1B2, IFI27, BTN3A1, SERTAD4, EPSTI1), which are a transcriptional response to SARS-CoV-2; 3) presenting interpretable functional effects of gene-gene interactions, gene-variants interactions using explicitly mathematical expressions; 4) presenting graphical tools for medical practitioners to understand the genomic signature patterns of the virus; 5) making suggestions on developing more efficient vaccines and antiviral drugs; 6) identifying potential genetic clues to other diseases due to COVID-19 infection. The remaining part of the paper is organized as follows. First, section 2 briefly reviews the studying methodology. Next, section 3 reports the data source, analysis results, and interpretations. Section 4 offers insights of addition twelve COVID-19 studies. Finally, Section 5 concludes the study.

## 2 Methodology

Many medical types of research, especially gene expression data related, applied the classical logistic regression as a starting base, then together with implementations of some advanced machine learning methods. However, Teng and Zhang (2021)^27^ point out that classical logistic regression can only model absolute treatments, not relative treatments. As a result, it has led (and will lead) to many supposedly efficient trials being wrongly concluded as inefficient. Four clinical trials, including one COVID-19 study trial, were illustrated in their paper. Their new AbRelaTEs regression model for medical data is much more advanced than the classical logistic regression as it greatly enhances interpretability and truly personalized medicine computability. Our new study in this paper is different from AbRelaTEs as we don’t deal with treatment and control, and we use a new innovative method to study the existence of functional effects of genes associated with SARS-CoV-2.

The competing risk factor classifier has been successfully applied in the literature^15;18;19;20;21^. This section briefly introduces necessary notations and formulas for self-contained due to different data structures used in this work. For continuous responses, the literature papers^28;29;30^ deal with max-linear computing factor models and max-linear regressions with penalization. The max-logistic classifier has some connections to the logistic polytomous models but with different structures^31;32;33^. This new innovative approach can be classified as either an AI algorithm or a machine learning algorithm. However, our new approach has an explicit formula and is interpretable.

Suppose *Y_i_* is the *i*th individual patient’s COVID-19 status (*y_i_* = 0 for COVID-19 free, *Y_i_* = 1 for infected) and 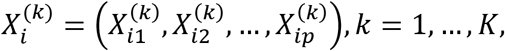 are the gene expression values with *p between* 15979 *and* 35784 genes in this study. Here *k* stands for the *k*th type of gene expression levels drawn based on *K* different biological sampling methodologies. Note that most published works set *K* = 1, and hence the superscript (*k*) can be dropped from the predictors. In this research paper, *K* = 4 as we have two datasets analyzed in Section 3, and in the first dataset, there are other ARIs patients with other viral infections or non-viral infections. Using a logit link (or any monotone link functions), we can model the risk probability 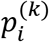 of the *i*th person’s infection status as:

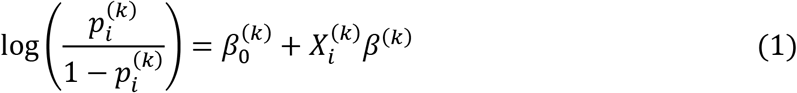

or alternatively, we write

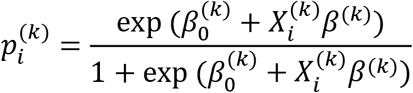

where 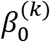 is an intercept, 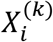 is a 1 × *p* observed vector, and *β*^(*k*)^ is a *p* × 1 coefficient vector which characterizes the contribution of each predictor (gene in this study) to the risk.

Considering there have been several variants of SARS-CoV-2 and multiple symptoms (subtypes) of COVID-19 diseases, it is natural to assume that the genomic structures of all subtypes can be different. Suppose that all subtypes of SARS-CoV-2 may be related to *G* groups of genes

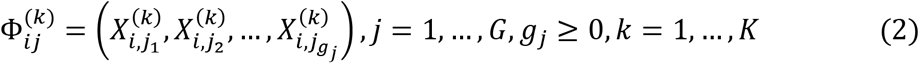

where *i* is the *i*th individual in the sample, *g_j_* is the number of genes in *j*th group.

The competing (risk) factor classifier is defined as

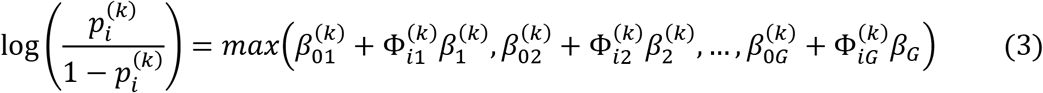

where 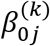’s are intercepts, 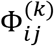 is a 1 × *g_j_* observed vector, 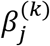 is a *g_j_* × 1 coefficient vector which characterizes the contribution of each predictor in the *j* group to the risk.

### Remark 1.

In (3), 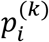 is mainly related to the largest component 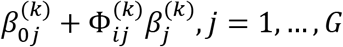, i.e., all components compete to take the most significant effect.

### Remark 2.

Taking 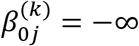, *j* = 2,…, *G*, (3) is reduced to the classical logistic regression, i.e., the classical logistic regression is a special case of the new classifier. Compared with black-box machine learning methods (e.g., random forest, deep learning (convolutional) neural networks (DNN, CNN)) and regression tree methods, each competing risk factor in (3) forms a clear, explicit, and interpretable signature with the selected genes. The number of factors corresponds to the number of signatures, i.e., *G*. This model can be a bridge between linear models and more advanced machine learning methods (black box) models. However, (3) remains the desired properties of interpretability, computability, predictability, and stability. Note that this remark is similar to Remark 1 in Zhang (2021)^19^.

We have to choose a threshold probability value to decide a patient’s class label in practice. Following the general trend in the literature, we set the threshold to be 0.5. As such, if 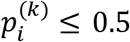, the *i*th individual is classified as disease-free; otherwise, the individual is classified as having the disease.

With the above-established notations and the idea of quotient correlation coefficient^34^, Zhang (2021)^19^ introduced a new machine learning classifier, smallest subset and smallest number of signatures (S4), for *K* = 1. We extend the S4 classifier from *K* = 1 to *K* = 4 as follows.

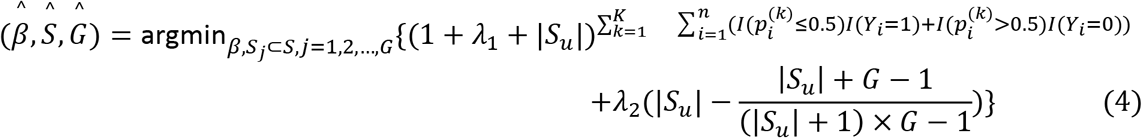

where *I*(·) is an indicative function, 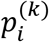 is defined in Equation (3), *S* = {1,2,…, 15979 *or* 35784} is the index set of all genes, *S_j_* = {*j*_*j*1_,…,*j_j,g_j__*}, *j* = 1,…, *G* are index sets corresponding to (2), *S_u_* is the union of {*S_j_, j* = 1,…, *G*}, |*S_U_*| is the number of elements in *S_u_, λ*_1_ ≥ 0 and *λ*_2_ ≥ 0 are penalty parameters, and 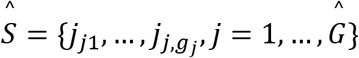 and 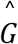> are the final gene set selected in the final classifiers and the number of final signatures.

### Remark 3.

The case of *K* = 1 corresponds to the classifier introduced in Zhang (2021)^19^. The case of *K* = 1and *λ_2_* = 0 corresponds to the classifier introduced in Zhang (2021)^15^.

## 3 Data Descriptions, Results and Interpretations

### 3.1 The data

Two COVID-19 datasets to be analyzed in this section are publicly available at https://github.com/czbiohub/covid19-transcriptomics-pathogenesis-diagnostics-results^22^ and as GSE152075^23^. The first dataset contains 15979 genes, 93 patients with NP/OP swaps PCR tested COVID-19 positive, 41 patients with viral acute respiratory illnesses (ARIs) and COVID-19 negative, and 100 non-viral acute respiratory illnesses (ARIs) COVID-19 negative. The second dataset contains 35784 genes, 430 individuals with NP/OP swaps PCR confirmed SARS-CoV-2 infection, and 54 negative controls. We note that many gene expression values in the second dataset are zero.

### 3.2 The competing factor classifiers and their resulting risk probabilities

Solving the optimization problem (4) among all genes (15979 and 35784), various competing classifiers can be identified with different combinations. As discussed in the introduction, the gene expression data used in this study were drawn from NP/OP swap PCR samples (not whole blood samples). Due to likely false positive and negative samples, 100% accurate classifiers with a single-digit number of genes do not exist. Also, with the same accuracy (smaller than 100%), different combinations of genes can be candidate classifiers. Therefore, we report the best-performed classifiers in this subsection. After an extensive Monte Carlo search of the best combinations of genes, five genes, ATP6V1B2, IFI27, BTN3A1 (Butyrophilin Subfamily 3 Member A1), SERTAD4 (SERTA Domain Containing 4), EPSTI1 (Epithelial Stromal Interaction 1), are found to form the S4 classifiers in Equation (4).

Given the first dataset has three categories (COVID-19 positive, ARIs with non-SARS-CoV-2 viral infection, ARIs without viral infection), we also study the classification between COVID-19 positive and ARIs with non-SARS-CoV-2 viral infection, and between COVID-19 positive and ARIs without viral infection, which leads to *K*=4 as stated in the prior subsection.

Note that in (3) each individual component itself is a classifier which has the following form

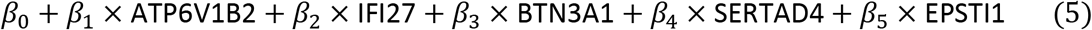

where (*β*_0_, *β*_1_,…, *β*_5_) are coefficients. In the subsequent subsections, we use tables to present individual (CF_*i,j*_) and combined (CFmax_*j*_) classifiers representing (5), where *i* is the index for a classifier, and *j* is for a dataset.

The risk probabilities of each component classifier are

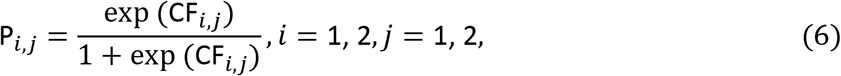

and the risk probabilities based on all three component classifiers together are

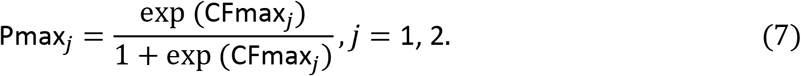

### 3.3 First dataset: Three-gene classifiers (*G*=1)

Note that the results in this subsection are not from our final best-performed classifiers. We found that a combination of ATP6V1B2 and IFI27 with many other genes can lead to high accuracy classifiers. We present their performance combined with the remaining genes of this paper’s best subset of five genes and one of the five critical genes found by Zhang^15^. Tables 1 and 2 summarize the results.

**Table 1.**
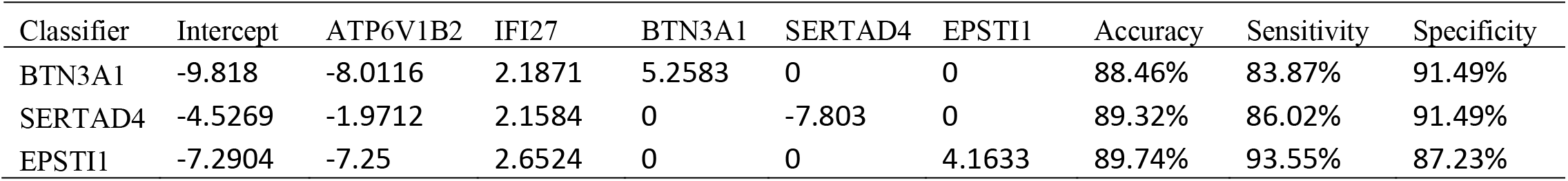
First dataset: Characteristics of the top performed individual genes together with ATP6V1B2 and IFI27 to form a three-gene classifier.

| Classifier | Intercept | ATP6V1B2 | IFI27 | BTN3A1 | SERTAD4 | EPSTI1 | Accuracy | Sensitivity | Specificity |
| --- | --- | --- | --- | --- | --- | --- | --- | --- | --- |
| BTN3A1 | -9.818 | -8.0116 | 2.1871 | 5.2583 | 0 | 0 | 88.46% | 83.87% | 91.49% |
| SERTAD4 | -4.5269 | -1.9712 | 2.1584 | 0 | -7.803 | 0 | 89.32% | 86.02% | 91.49% |
| EPSTI1 | -7.2904 | -7.25 | 2.6524 | 0 | 0 | 4.1633 | 89.74% | 93.55% | 87.23% |

**Table 2.** First dataset: Characteristics of RIPK3 together with ATP6V1B2 and IFI27.

| Classifier | Intercept | ATP6V1B2 | IFI27 | RIPK3 | Accuracy | Sensitivity | Specificity |
| --- | --- | --- | --- | --- | --- | --- | --- |
| RIPK3 | -1.2487 | -5.7586 | 1.3916 | 9.902 | 87.2% | 76.3% | 94.3% |

Tables 1 and 2 show that the coefficient signs of ATP6V1B2 and IFI27 are the same across all individual classifiers, which is a strong indication that they are truly associated with the virus. Although gene RIPK3 plays a key role in the perfect classifier identified in Zhang^15^, its performance is inferior to the other three genes identified from NP/OP swap PCR samples in this paper. This phenomenon reflects the discussions in the Introduction that RIPK3 is related to the natural essence of COVID-19, while ATP6V1B2, IFI27, BTN3A1, SERTAD4, and EPSTI1 contain more information about SARS-CoV-2.

We note that BTN3A1 combinations with ATP6V1B2 and IFI27 can have numerous types, which also leads to the same accuracy; for SERTAD4, there are numerous combinations with ATP6V1B2 and IFI27; and the same is true for EPSTI1. The coefficients listed in Table 1 are just a particular type of coefficient. Also, for EPSTI1, we can get different sensitivities and specificities while maintaining the same accuracy. Among four genes (BTN3A1, SERTAD4, EPSTI1, and RIPK3), EPSTI1 performs best in Tables 1 and 2. This empirical evidence proves that ATP6V1B2 and IFI27 are at the center of genes associated with SARS-CoV-2.

### 3.4 First dataset: Five-gene classifiers and the existence of variants

Our extensive Monte Carlo search leads to the best solution of the accuracy of 91.82% to the optimization problem (4) by five genes, i.e., ATP6V1B2, IFI27, BTN3A1, SERTAD4, and EPSTI1 though the solution is not unique. After comparing solutions for all three categories in the first dataset, these five genes stand out. Tables 3–5 summarize the results.

**Table 3.** First dataset: Characteristics of the top performed five-gene classifier. CF1 and CF2 are the first and second individual classifiers for data COVID-19 patients vs. other viral ARIs and non-viral infection patients.

| Classifier | Intercept | ATP6V1B2 | IFI27 | BTN3A1 | SERTAD4 | EPSTI1 | Accuracy | Sensitivity | Specificity |
| --- | --- | --- | --- | --- | --- | --- | --- | --- | --- |
| CF1 | 9.193 | -1.8935 | 1.5774 | 0 | -4.3303 | 0 | 87.61% | 81.72% | 91.49% |
| CF2 | -7.2786 | -5.2993 | 0 | 3.2572 | 0 | 2.34 | 86.32% | 76.34% | 92.91% |
| max{CF1, CF2} |  |  |  |  |  |  | 91.88% | 94.62% | 90.07% |

**Table 4.** First dataset: Characteristics of the top performed five-gene classifier. CF1 and CF2 are the first and second individual classifiers for data COVID-19 patients vs. other viral infection ARIs but non-viral infection patients.

| Classifier | Intercept | ATP6V1B2 | IFI27 | BTN3A1 | SERTAD4 | EPSTI1 | Accuracy | Sensitivity | Specificity |
| --- | --- | --- | --- | --- | --- | --- | --- | --- | --- |
| CF1 | -2.052 | 0 | 3.9086 | 2.5578 | 0 | -9.6586 | 70.15% | 62.37% | 87.8% |
| CF2 | 5.5979 | -7.4352 | 0 | 0 | 8.3704 | 4.4936 | 76.12% | 74.19% | 80.49% |
| max{CF1, CF2} |  |  |  |  |  |  | 91.04% | 97.85% | 75.61% |

**Table 5.** First dataset: Characteristics of the top performed five-gene classifier. CF1 and CF2 are the first and second individual classifiers for data COVID-19 patients vs. non-viral infection ARIs patients.

| Classifier | Intercept | ATP6V1B2 | IFI27 | BTN3A1 | SERTAD4 | EPSTI1 | Accuracy | Sensitivity | Specificity |
| --- | --- | --- | --- | --- | --- | --- | --- | --- | --- |
| CF1 | -2.2381 | -7.9733 | 0 | 4.5448 | 0 | 4.7567 | 90.16% | 81.72% | 98% |
| CF2 | -2.1003 | -4.8036 | 4.0849 | 0 | -9.9738 | 0 | 90.16% | 82.8% | 97% |
| max{CF1, CF2} |  |  |  |  |  |  | 96.37% | 95.70% | 97% |

In section 3.3, we forced ATP6V1B2 and IFI27 to be members in each classifier, while the best performance classifiers in this section reveal they can function separately, which tells that a gene’s function heavily depends on other genes’ function, i.e., gene-gene interactions, and genedisease subtype interactions. Furthermore, such a phenomenon suggests SARS-CoV-2 variants/subtypes are heterogeneous. As a result, models without differentiating gene-gene interactions and gene-variants interactions can be suboptimal.

Table 6 demonstrates part of patients’ expression values of the five critical genes, competing classifier factors, and predicted probabilities. Note that due to relatively very large scales in Columns CF-1, CF-2, CFmax, they are rescaled by a division of 100 when computing the risk probabilities as very large values can result in an overflow in computation. The validity of rescaling was justified in Zhang^17^.

**Table 6.** First dataset: Expression values of the five critical genes, competing classifier factors, predicted probabilities.

| #ID | Status | ATP6V1B2 | IFI27 | BTN3A1 | SERTAD4 | EPSTI1 | CF1 | CF2 | CFmax<sub>1</sub> | P<sub>1</sub> | P<sub>2</sub> | P-1 |
| --- | --- | --- | --- | --- | --- | --- | --- | --- | --- | --- | --- | --- |
| e-202 | 0 | 277 | 604 | 104 | 158 | 138 | -246.7 | -813.52 | -246.7 | 0.08 | 0.00 | 0.08 |
| e-080 | 0 | 866 | 103 | 82 | 76 | 94 | -1797.2 | -4109.42 | -1797.2 | 0.00 | 0.00 | 0.00 |
| e-287 | 0 | 3127 | 717 | 271 | 233 | 151 | -5789.8 | -15342.2 | -5789.8 | 0.00 | 0.00 | 0.00 |
| e-753 | 1 | 1053 | 2029 | 766 | 214 | 819 | 289.2 | -1176.0 | 289.2 | 0.95 | 0.00 | 0.95 |
| e-751 | 1 | 253 | 1423 | 266 | 114 | 369 | 1281.1 | 381.87 | 1281.1 | 1.00 | 0.98 | 1.00 |
| e-520 | 0 | 617 | 344 | 120 | 11 | 559 | -664.1 | -1578.0 | -664.1 | 0.00 | 0.00 | 0.00 |
| e-505 | 0 | 721 | 240 | 298 | 10 | 500 | -1020.8 | -1687.4 | -1020.8 | 0.00 | 0.00 | 0.00 |
| i-083 | 0 | 191 | 320 | 119 | 72 | 71 | -159.5 | -465.7 | -159.5 | 0.17 | 0.01 | 0.17 |
| e-764 | 0 | 1667 | 202 | 76 | 3 | 1232 | -2841.6 | -5710.8 | -2841.6 | 0.00 | 0.00 | 0.00 |
| e-451 | 0 | 1880 | 24 | 98 | 2 | 27 | -3521.4 | -9587.6 | -3521.4 | 0.00 | 0.00 | 0.00 |
| e-285 | 0 | 794 | 826 | 530 | 392 | 300 | -1888.8 | -1786.6 | -1786.6 | 0.00 | 0.00 | 0.00 |
| e-254 | 0 | 512 | 253 | 195 | 388 | 69 | -2241.4 | -1923.9 | -1923.9 | 0.00 | 0.00 | 0.00 |
| e-726 | 1 | 398 | 1395 | 362 | 96 | 567 | 1040.3 | 389.5 | 1040.3 | 1.00 | 0.98 | 1.00 |

Figure 1 presents critical gene expression levels and risk probabilities corresponding to different combinations in the first dataset and Tables 3–5. It can be seen that each plot shows a genomic signature pattern and functional effects of the genes involved.

**Figure 1:**
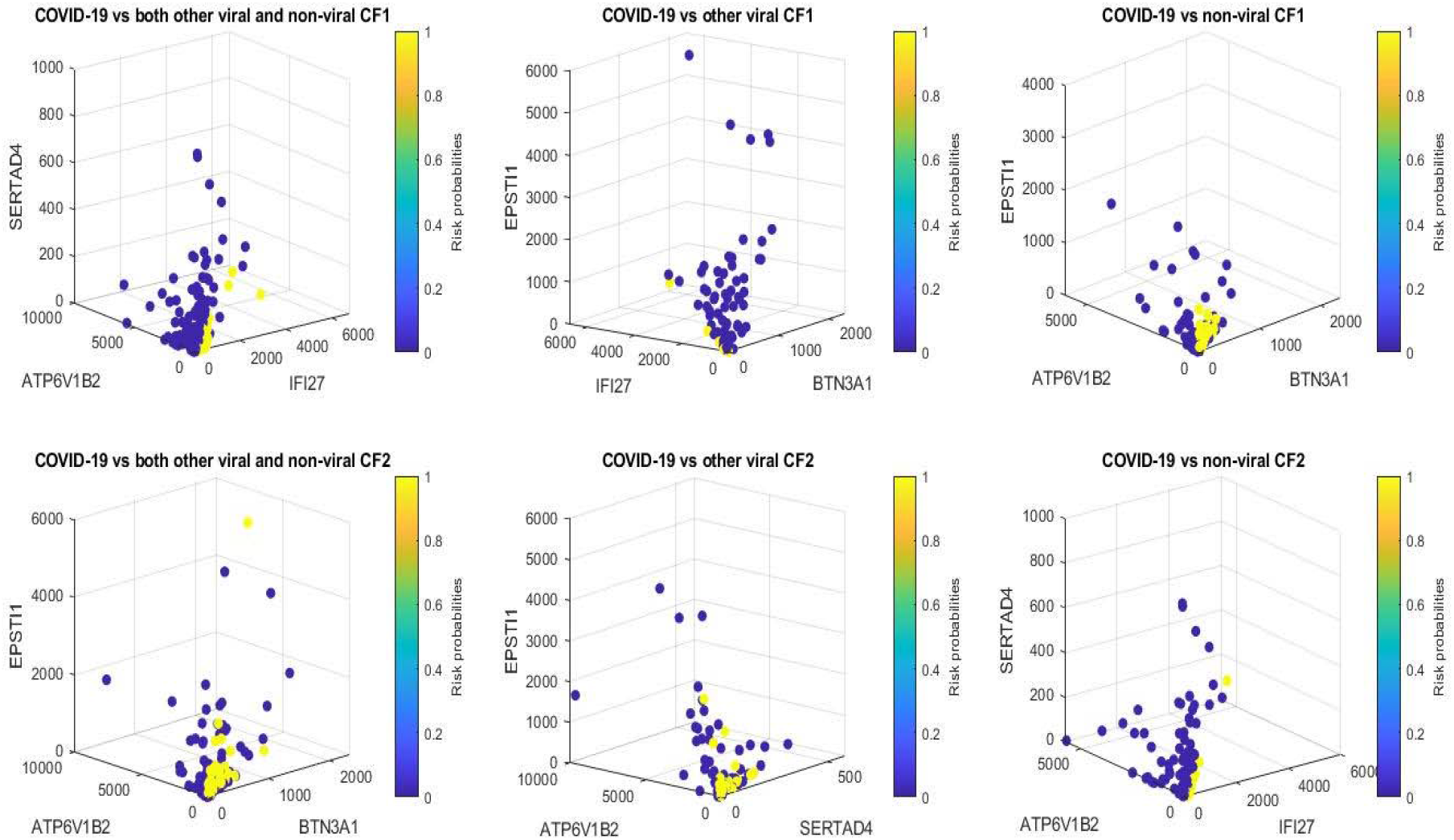
COVID-19 classifiers in Tables 3–5: Visualization of gene-gene relationship and gene-risk probabilities. Note that 0.5 is the probability threshold.

From Tables 1–5, we can immediately see that the coefficient signs associated with ATP6V1B2 are uniformly negative, which shows that increasing the expression level of ATP6V1B2 will decrease the virus (SARS-CoV-2) strength; the coefficient signs associated with IFI27 are uniformly positive, which shows that decreasing the expression level of IFI27 will decrease the virus (SARS-CoV-2) infection strength. Such functional effects of ATP6V1B2 and IFI27 can also be clearly seen in Figure 1 around origins which show the higher the IFI27 level, the higher the risk probability (yellow color); the higher the ATP6V1B2 level, the lower the risk probability (blue color). These observations show that ATP6V1B2 and IFI27 are in the circle of genes associated with SARS-COV-2. BTN3A1 appears three times in Tables 3–5 with positive coefficients, which shows decreasing the expression level of BTN3A1 will decrease the virus (SARS-CoV-2) infection strength. The coefficient signs of SERTAD4 and the coefficient signs of EPSTI1 show both positive and negative in Tables 3–5 depending on how genes are combined. These phenomena explain the reason SARS-CoV-2 variants have emerged, as variants can be related to different coefficient signs corresponding to genes.

Figure 2 is a Venn diagram illustrating each classifier’s performance and the combined classifier. In Venn diagram, those patients who fall in the intersections are relatively easy to be tested and confirmed positive, while for those who only fall in one category, it is relatively hard to test and confirm their status. Two individual classifiers can be explained as having two times COVID-19 tests using two different testing procedures (kits), and with both tests being positive, the probability of infection will be higher depending on the sensitivity and the specificity of each test. Summarizing Tables 3–5 and Figure 2, mathematically speaking, SARS-CoV-2 can have 3 × 3 × 3 × 4 = 108 variants, with some of them being insignificant from dominant ones while some of them being dominant and having emerged (or will emerge), where the multiplier 3 corresponds to 3 classes in one Venn diagram, and similarly, other numbers are interpreted. Such many variants may offer a genomic clue to what has been found in Chertow et al. (2021)^26^. We note that the joint functional effects of genes are not directly observable, and the meaning of variants is defined by the joint functional effects. As a result, the variants of the virus are not directly referred to as what has been known in the literature and practice.

**Figure 2:**
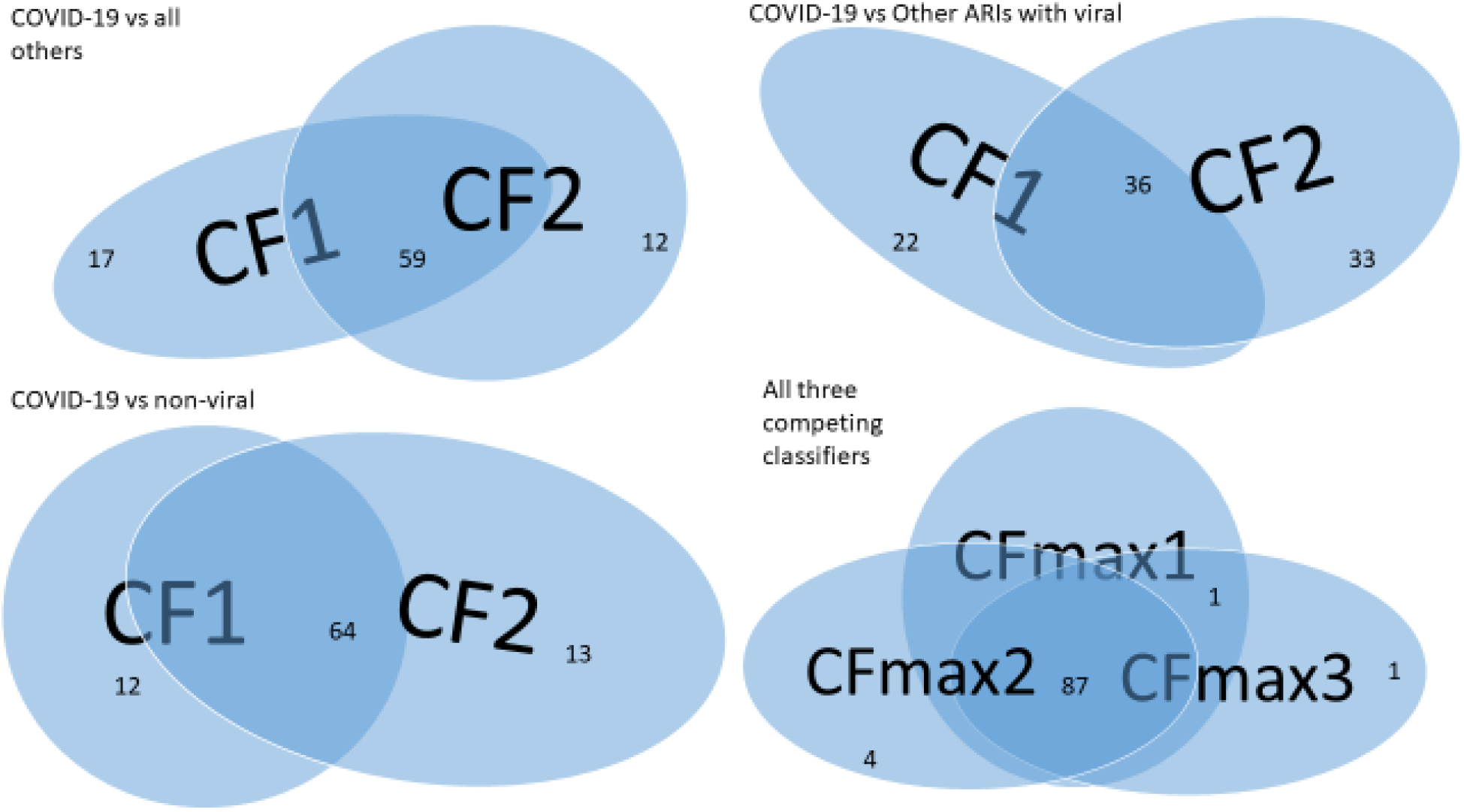
Venn diagram of variants of SARS-CoV-2 (the first dataset): Top-left panel is for COVID-19 vs. all others; Top-right panel is for COVID-19 vs. other viral; Bottom-left panel is for COVID-19 vs. non-viral; Bottom-right panel is for all three together.

Comparing the individual classifiers and combined classifiers among COVID-19 vs. all others, COVID-19 vs. ARIs with other viral infection, and COVID-19 vs. without viral infection, we see that the combined classifier for the case of COVID-19 vs. without viral infection works the best. We found some ARIs with other viral infections may be COVID-19 patients but not yet confirmed. Applying the classifier in Figure 2 bottom-right panel can get sensitivity up to 98.94% with a slight loss of specificity.

The five genes, ATP6V1B2, IFI27, BTN3A1, SERTAD4, and EPSTI1, performed better in classifying patients in their respective groups in the first dataset. Therefore, a natural question will be whether or not the accuracies are overestimated. Next, we address this question in two aspects.

In the literature, in order to avoid overfitting data, cross-validation (CV) has been widely utilized in model building and inference. However, this methodology is only working when samples are drawn from a homogeneous population. When samples are from heterogeneous populations, CV methods will lead to inaccurate classification results, and eventually, the results are not interpretable. Having observed COVID-19 disease subtypes and SARS-CoV-2 variants, heterogeneous populations of all genes are the basic structure of COVID-19 genomics (transcriptional data). As a result, the classical CV method is not applicable in our studies.

Alternatively, given the fundamental task is to identify critical genes and their joint effects as high-performance genomic biomarkers, we can directly fit the genes identified from the first dataset to several other datasets to test the fitted models and their prediction accuracy. We adopt this approach in this paper.

Additionally, using the existing methods to identify high-performance genes, dozens have been reported in the literature with lower accuracy than the single-digit number of genes in our new work. If we conclude that the genes identified in this study are overestimated, then we argue that the gene sets with doubled or even tripled numbers of genes should definitely be overestimated and must be useless or not meaningful at all. Therefore, all biological inferences based on those double/tripled numbers of genes can be misleading.

### 3.5 Second dataset: Five-gene classifiers and the existence of variants

In this subsection, we test their performance in a second dataset. One significant difference between these two datasets is that the patients in the first study (dataset) are either COVID-19 positive or ARIs with other viral infection or ARIs without viral infection, while the patients in the second study (dataset) are NP/OP swap PCR confirmed SARS-CoV-2 infection or negative controls. As a result, genes found to be critical from the first dataset can be thought of as SARS-CoV-2 specific. It turned out that those five genes are also the best subset for the second dataset. Table 7 presents the results from an individual classifier. Data are ln(raw+1) normalized.

**Table 7.**
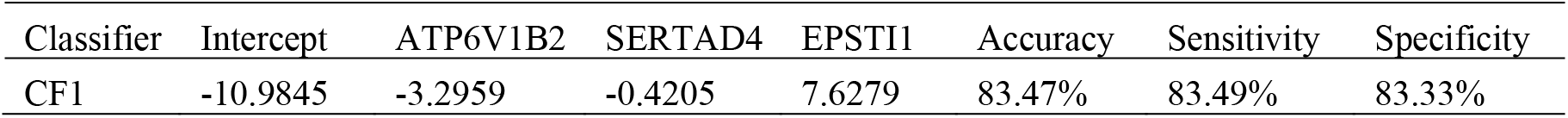
Second dataset: Characteristics of the top performed five-gene classifier. CF1 stands for the first individual classifier for data COVID-19 positive vs. COVID-19 negative.

| Classifier | Intercept | ATP6V1B2 | SERTAD4 | EPSTI1 | Accuracy | Sensitivity | Specificity |
| --- | --- | --- | --- | --- | --- | --- | --- |
| CF1 | -10.9845 | -3.2959 | -0.4205 | 7.6279 | 83.47% | 83.49% | 83.33% |

We can see that the signs of ATP6V1B2, SERTAD4 and EPSTI1 in CF1 remain the same as their counterparts in Tables 1–5. This table again supports our earlier claim that ATP6V1B2, IFI27 are in the circle of critical genes associated with SARS-CoV-2. Table 7 also reveals that the information derived using the key genes derived from other datasets can be weak due to weak data quality (e.g., very noisy, no signals). On the other hand, our method can still perform satisfactorily with an overall accuracy of 83.47, sensitivity of 83.49%, and specificity of 83.33%, proving the importance of the identified critical genes and showing the new method’s superiority.

Note that the individual classifier CF1 in the second dataset has a different combination compared with the counterparts in the first dataset. This phenomenon can be explained by the different patients’ attributes from these two datasets. Next, we compute the correlations among those five genes for each dataset. Table 8 presents pairwise correlations in a matrix form in which the upper triangle is for the first dataset, and the lower triangle is for the second dataset.

**Table 8.** Pairwise correlation coefficients: The upper triangle is for the first dataset, and the lower triangle is for the second dataset.

|  | ATP6V1B2 | IFI27 | BTN3A1 | SERTAD4 | EPSTI1 |
| --- | --- | --- | --- | --- | --- |
| ATP6V1B2 | 1 | 0.208 | 0.5416 | 0.051 | 0.5415 |
| IFI27 | 0.4031 | 1 | 0.5463 | 0.3084 | 0.5616 |
| BTN3A1 | 0.69 | 0.3823 | 1 | 0.25 | 0.7527 |
| SERTAD4 | 0.3417 | 0.3302 | 0.2663 | 1 | 0.0079 |
| EPSTI1 | 0.6531 | 0.3366 | 0.6562 | 0.1791 | 1 |

Table 8 shows different correlation structures among the five genes, which makes the difference in classifiers between the two datasets reasonable.

## 4 Genomic differences between NP/OP swaps PCR samples and whole blood samples

In this section, we use additional twelve datasets to cross-validate the genes identified in Section 3. These datasets include GSE152641^37^, GSE155454^38^, GSE163151^39^, GSE166190^40^, GSE166253^41^, GSE166530^42^, GSE177477^43^, GSE179448^44^, GSE184401^45^, GSE189039^46;47^, GSE190680^48^, and GSE201530^47;49^.

We first used GSE152641 and GSE166530 to form a combined dataset to empirically justify the genes identified in Section 3 and those genes (ABCB6, KIAA1614, MND1, SMG1, RIPK3, CDC6, ZNF282, CEP72) published in our earlier work^17^ are functionally distinct in SARS-CoV-2 and COVID-19. GSE152641 has the overall design being total RNA sequencing from whole blood of 62 COVID-19 patients and 24 healthy controls, the platform being GPL24676 illumina NovaSeq 6000 (Homo sapiens), and genome build being GRCh38. GSE166530 has the overall design of nasopharyngeal or oropharyngeal swabs PCR samples with 36 COVID-19 positives and 5 negatives. Its platform and genome build are the same as those of GSE152641. We combine the 62 COVID-19 whole blood sampled patients from GSE152641 and 36 COVID-19 positive NP/OP swap samples together to form a new dataset. Figure 3 plots expression levels (raw counts) of the new dataset.

**Figure 3.**
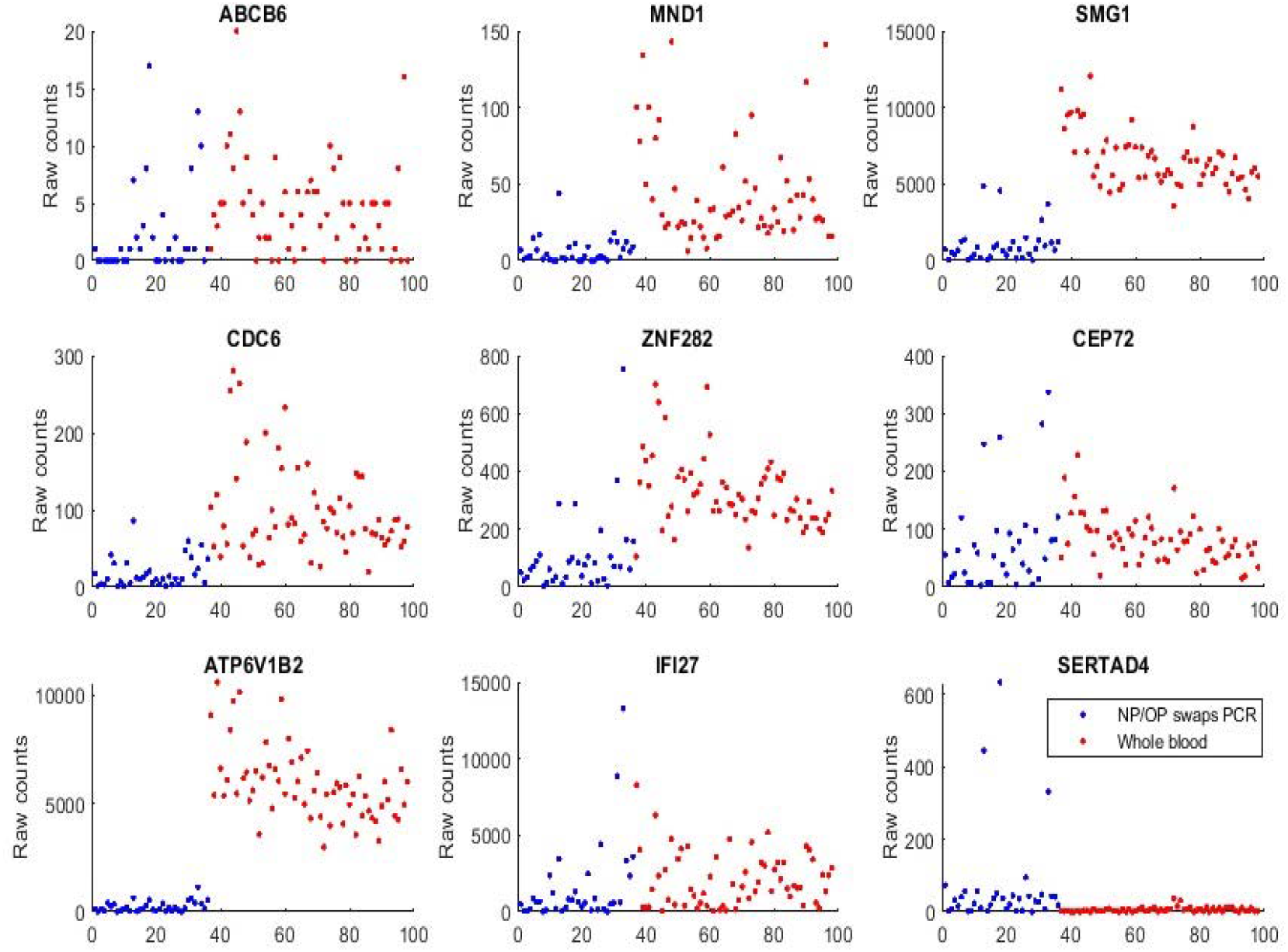
Gene expression raw counts from COVID-19 positives. The red-colored dots represent patients from GSE152641 whole blood samples. The blue dots represent patients from GSE166530 NP/OP swap PCR samples.

We can see that samples from both populations show some similarities in expression level ranges with ABCB6, CEP72, and IFI27, which justifies the graphical comparison being feasible since GSE1552641 and GSE166530 have some subtle differences in their data generating processes though they use the same platform and genomic build.

We can see that ATP6V1B2 shows a completely separable pattern between the two populations. MND1, SMG1, CDC6, ZNF282 all have higher expression levels in the whole blood than in NP/OP swaps.

We found that SERTAD4’s transcriptomic data in whole blood samples are almost all zeros or very small in Figure 3 (GSE1552641) and other whole blood samples to be analyzed. This phenomenon tells that SERTAD4 is a phenomenon of symptom.

Analyzing GSE152641 separately, we obtain the following Table 9.

**Table 9.** GSE152641: Characteristics of the top performed four-gene classifier. CF1 and CF2 are the first and second individual classifiers for data COVID-19 patients vs. healthy control.

| Classifier | Intercept | KIAA1614 | RIPK3 | CDC6 | ZNF282 | Accuracy | Sensitivity | Specificity |
| --- | --- | --- | --- | --- | --- | --- | --- | --- |
| CF1 | -5.7093 |  | 8.4656 | 5.8485 | -9.3695 | 80.23% | 72.58% | 100% |
| CF2 | -6.8734 | 5.9693 | -2.9708 | 8.6925 |  | 77.91% | 69.35% | 100% |
| Max{CF1,CF2} |  |  |  |  |  | 98.84% | 98.39% | 100% |

Comparing Table 9 and our earlier results^17^, we see that the combination of CDC6 and ZNF282 is extended to RIPK3 and KIAA1614, which suggests CDC6 and ZNF282 can be core genes and other genes, e.g., CEP72, RIPK3 and KIAA1614 can be substituted.

GSE155454 has an overall design: RNA was extracted from whole blood collected from 27 COVID-19 patients from the Singapore cohort after retrospective matching and 6 healthy controls. Timepoints selected for extraction were during active infection (PCR-positive, median 8 days PIO) and recovered (PCR-negative; median 21 days PIO). The platform was GPL20301 Illumina HiSeq 4000 (Homo sapiens). Table 10 presents our classification results based on the genes identified in Section 3.

**Table 10.** GSE155454: Characteristics of the top performed three-gene classifier CF1 for data COVID-19 positive vs. negative and healthy control.

| Classifier | Intercept | RIPK3 | ZNF282 | IFI27 | Accuracy | Sensitivity | Specificity |
| --- | --- | --- | --- | --- | --- | --- | --- |
| CF1 | -7.9946 | -7.1725 | 6.9103 | 5.7519 | 93.75% | 89.66% | 84.62% |

We note that this data collection included COVID-19 recovered, i.e., COVID-19 negative. The coefficient signs of ZNF282 and IFI27 obviously differ from our earlier work^17^ and in Table 6. One possible reason is that the recovered patient has different gene expression levels compared with their COVID-19 naïve counterparts, i.e., SARS-CoV-2 infection effects at the genomic level weren’t completely faded away. Nevertheless, CF1 in Table 10 still leads to high-performance 93.75% accuracy.

GSE163151 conducted RNA sequencing (RNA Seq) to analyze nasopharyngeal (NP) swab and whole blood (WB) samples from 333 COVID-19 patients and controls, including patients with other viral and bacterial infections. The platform was GPL24676 Illumina NovaSeq 6000 (Homo sapiens). We take a subset of the data to study the genes identified in Section 3 and our earlier work. In particular, 138 NP swap samples and 7 whole blood samples were used. Table 11 presents our classification results.

**Table 11.**
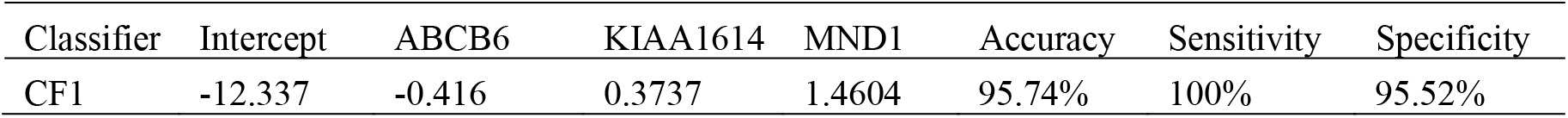
GSE163151: Characteristics of the top performed three-gene classifier CF1 for data whole blood vs. NP/OP swaps.

| Classifier | Intercept | ABCB6 | KIAA1614 | MND1 | Accuracy | Sensitivity | Specificity |
| --- | --- | --- | --- | --- | --- | --- | --- |
| CF1 | -12.337 | -0.416 | 0.3737 | 1.4604 | 95.74% | 100% | 95.52% |

With the accuracy of 95.74%, clearly, we see that COVID-19 NP swap samples and whole blood samples have different gene-gene interactions among those critical genes identified in Section 3 and our earlier work^17^. Therefore, scientists should pay attention to this dissimilarity, which is fundamental to fighting against the COVID-19 pandemic.

GSE166190’s overall design is a transcriptomic analysis of whole blood from SARS-COV-2 infected participants and their SARS-CoV-2 negative household contacts. In the analysis, transcriptomic data of an individual were collected in 5-time intervals according to the calculated days POS: interval 1 (0-5), interval 2 (6-14), interval 3 (15-22), interval 4 (23-35), and interval 5 (36-81). The platform was GPL20301 Illumina HiSeq 4000 (Homo sapiens). Table 12 presents our analysis of the data.

**Table 12.** GSE166190: Characteristics of the top performed six-gene classifier. CF1, CF2, CF3 are the first, second and third individual classifiers for data on COVID-19 patients vs. healthy control. The data were natural logarithm transformed as ln(KIAA1614/10+1), ln(MND1+1), ln(RIPK3/10+1), ln(SMG1/100+1), ln(ZNF282/10+1), ln(CEP72+1).

| Classifier | Intercept | KIAA1614 | MND1 | RIPK3 | SMG1 | ZNF282 | CEP72 | Accuracy | Sensitivity | Specificity |
| --- | --- | --- | --- | --- | --- | --- | --- | --- | --- | --- |
| CF1 | 25.7352 |  |  | 11.4885 | -16.3554 | -1.6889 |  | 31.63% | 19.28% | 100% |
| CF2 | 8.5694 | -9.6995 | 4.0413 |  |  |  | 2.342 | 62.24% | 55.42% | 100% |
| CF3 | -12.6727 | -5.3787 |  | 11.2971 |  |  | -3.1795 | 27.55% | 14.46% | 100% |
| CF max |  |  |  |  |  |  |  | 77.55% | 73.49% | 100% |

Different from GSE155454, this study’s time intervals are quite wide. We used six critical genes identified in our earlier work^17^ to reach a 77.55% accuracy which is much lower than our other analysis in the COVID-19 study though it is already an accepted rate. A possible reason is that in this data, gene-gene interactions from the interval 1 (0-5days) to the follow-up intervals are different, which decreased the sensitivities of our CFi classifiers. However, we obtained 100% specificity with all individuals being tested up to five times. In our Supplementary full data table, we found that we had interval 1 being 100% sensitivity and some of interval 2 being 100% sensitivity. As such, it may be safe to say that the genes in our earlier work^17^ worked perfectly.

GSE166253 studied transcriptomic characteristics and impaired immune function of patients who retested positive (RTP) for SARS-CoV-2 RNA. The platform was GPL20795 HiSeq X Ten (Homo sapiens). The data contains 10 retested positive patients, 6 convalescent patients, and 10 healthy controls who were enrolled for analysis of the immunological characteristics of their peripheral blood mononuclear cells. Table 13 reports our fitting results.

**Table 13.**
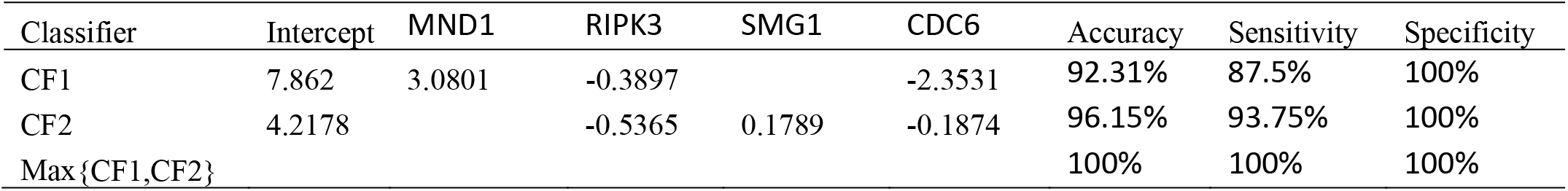
GSE166253: Characteristics of the top performed four-gene classifier. CF1 and CF2 are the first and second individual classifiers for data COVID-19 patients vs. healthy control.

| Classifier | Intercept | MND1 | RIPK3 | SMG1 | CDC6 | Accuracy | Sensitivity | Specificity |
| --- | --- | --- | --- | --- | --- | --- | --- | --- |
| CF1 | 7.862 | 3.0801 | -0.3897 |  | -2.3531 | 92.31% | 87.5% | 100% |
| CF2 | 4.2178 |  | -0.5365 | 0.1789 | -0.1874 | 96.15% | 93.75% | 100% |
| Max{CF1,CF2} |  |  |  |  |  | 100% | 100% | 100% |

The table shows that the gene-gene interactions are different among RTP and convalescent patients. It is interesting to note that we had 100% accuracy in this data analysis.

GSE166530 was used in Figure 3 with its COVID-19 positive patients’ NP swap sampled gene expression levels. In addition, we conducted a separate classification analysis using the five genes identified in Section 3. Table 14 reports the results.

**Table 14.** GSE166530: Characteristics of the top performed five-gene classifier. CF1 and CF2 are the first and second individual classifiers for data COVID-19 patients vs. healthy control

| Classifier | Intercept | ATP6V1B2 | IFI27 | BTN3A1 | SERTAD4 | EPSTI1 | Accuracy | Sensitivity | Specificity |
| --- | --- | --- | --- | --- | --- | --- | --- | --- | --- |
| CF1 | -11.1266 | 8.5087 | -1.4154 |  |  | -7.6515 | 31.71% | 22.22% | 100% |
| CF2 | 6.8238 | 0.4763 |  |  | -1.9013 | 0.2038 | 86.11% | 86.11% | 100% |
| max{CF1, CF2} |  |  |  |  |  |  | 95.12% | 94.44% | 100% |

This table shows different coefficient patterns different from Table 7. We note that we only have 5 healthy individuals in control. Interestingly, if we use the five genes identified in our earlier work^15;17^, we can get 100% accuracy. This Indian cohort is worth further looking into its genegene and subvariant interactions. However, we didn’t find additional characteristics available to study.

GSE177477 is a Pakistan cohort study. Its overall design is that COVID-19 cases with positive respiratory samples SARS-CoV-2 and healthy Controls cases were recruited. Blood transcriptomes were analyzed using Clariom S RNA Microarray, Affymetrix Inc. The platform was GPL23159 [Clariom_S_Human] Affymetrix Clariom S Assay, Human (Includes Pico Assay). We used 11 symptomatic samples and 18 healthy control samples to test our earlier work identified genes’ predicting accuracy. We obtained 100% accuracy in this analysis. The results are presented in Table 15.

**Table 15.** GSE177477: Characteristics of the top performed four-gene classifier. CF1 and CF2 are the first and second individual classifiers for data COVID-19 patients vs. healthy control.

| Classifier | Intercept | SMG1 | CDC6 | ZNF282 | CEP72 | Accuracy | Sensitivity | Specificity |
| --- | --- | --- | --- | --- | --- | --- | --- | --- |
| CF1 | 0.6104 | 2.3501 | 1.6222 |  | -8.9165 | 79.31% | 45.45% | 100% |
| CF2 | -2.0531 | -0.5364 | 13.4123 | -10.0557 |  | 93.10% | 81.82% | 100% |
| Max{CF1,CF2} |  |  |  |  |  | 100% | 100% | 100% |

The coefficient signs of CDC6, ZNF282, CEP71 are consistent with our earlier work^17^. Again, this study leads to the importance of CDC6 and ZNF282.

GSE179448 conducted RNAseq analysis of human CD4+ regulatory Tregs and Tconvs in COVID-19 and healthy donors isolated from peripheral blood. We used 22 hospitalized COVID-19 samples and 15 healthy control samples to test our earlier work identified genes’ predicting accuracy. The results are presented in Table 16.

**Table 16.** GSE179448: Characteristics of the top performed five-gene classifier. CF1 and CF2 are the first and second individual classifiers for data on COVID-19 hospitalized vs healthy control

| Classifier | Intercept | KIAA1614 | MND1 | RIPK3 | CDC6 | CEP72 | Accuracy | Sensitivity | Specificity |
| --- | --- | --- | --- | --- | --- | --- | --- | --- | --- |
| CF1 | 4.328 |  | -1.5254 |  | 8.7869 | -4.5027 | 81.08% | 72.72% | 93.33% |
| CF2 | 10.0917 | 7.9273 |  | -4.3736 |  | 6.7933 | 48.65% | 13.64% | 100% |
| max{CF1, CF2} |  |  |  |  |  |  | 89.19% | 86.36% | 93.33% |

We obtained an 89.19% overall accuracy in this study. One possible reason may be that the platform is GPL18573 Illumina NextSeq 500 (Homo sapiens), compared with GPL24676 illumina NovaSeq 6000 (Homo sapiens) which led to higher accuracy.

GSE184401 used a platform of GPL24676--Illumina NovaSeq 6000 (Homo sapiens). Its overall design is a RNA-seq analysis in the peripheral blood mononuclear cell isolated shortly from the initial infection. All individuals were COVID-19 confirmed with three types, severe condition with the secondary infection, severe condition without secondary infection, and mild infection. We present our results of four genes from our earlier work^17^ and three from Section 3 in Table 17.

**Table 17.** GSE184401: Characteristics of the top performed seven-gene classifier. CF1, CF2, CF3 are the first, second and third individual classifiers for data COVID-19 severe condition vs. mild infection.

| Classifier | Intercept | KIAA1614 | CDC6 | ZNF282 | CEP72 | ATP6V1B2 | IFI27 | BTN3A1 | Accuracy | Sensitivity | Specificity |
| --- | --- | --- | --- | --- | --- | --- | --- | --- | --- | --- | --- |
| CF1 | -1.4488 |  | 6.9615 |  | -2.125 |  | -3.828 |  | 76.74% | 59.09% | 95.24% |
| CF2 | 10.8726 | 9.3158 |  |  | -4.309 |  | -0.19 |  | 69.77% | 40.91% | 100% |
| CF3 | 5.1235 |  |  | 6.758 |  | 2.5267 |  | -4.0934 | 88.37% | 77.27% | 100% |
| CF max |  |  |  |  |  |  |  |  | 95.35% | 95.45% | 95.24% |

From this analysis, we see that gene-gene interactions are different after infection with different severe conditions.

GSE189039 has the overall design as RNA-seq was performed with peripheral blood mononuclear cells (PBMCs) of COVID-19 patients infected by SARS-CoV-2 Beta variant (Beta) and SARS-CoV-2 naïve vaccinated individuals. The platform was GPL24676 Illumina NovaSeq 6000 (Homo sapiens). Our analysis results are presented in Table 18.

**Table 18.** GSE189039: Characteristics of the top performed three-gene classifier CF1 for data COVID-19 vs. healthy control.

| Classifier | Intercept | ABCB6 | MND1 | CEP72 | Accuracy | Sensitivity | Specificity |
| --- | --- | --- | --- | --- | --- | --- | --- |
| CF1 | 4.742 | -0.001 | 0.0402 | -0.072 | 100% | 100% | 100% |

It is interesting to point out that we used only one classifier CF1 to reach 100% accuracy.

GSE190680 has an overall design: RNA-seq was performed with peripheral blood mononuclear cells (PBMCs) of COVID-19 patients infected by SARS-CoV-2 Alpha variant with or without the escape mutation. The platform was GPL24676 Illumina NovaSeq 6000 (Homo sapiens). Note that all patients in this study were COVID-19 patients infected by SARS-CoV-2 Alpha variant. We used our identified critical genes to test the ability to separate E484K escape mutation. Table 19 presents the results.

**Table 19.** GSE190680: Characteristics of the top performed three-gene classifier CF1 for data COVID-19 vs. healthy control.

| Classifier | Intercept | ABCB6 | CDC6 | CEP72 | Accuracy | Sensitivity | Specificity |
| --- | --- | --- | --- | --- | --- | --- | --- |
| CF1 | 9.8031 | -2.1852 | 1.4385 | 3.1508 | 84% | 76.67% | 87.14% |

With an overall accuracy of 84%, it is safe to say that the three genes ABCB6, CDC6, CEP72 have the ability to predict E484K escape mutation.

In GSE201523, RNA-seq was performed with peripheral blood mononuclear cells (PBMCs) of COVID-19 patients infected by SARS-CoV-2 Omicron variant. The platform was GPL24676 Illumina NovaSeq 6000 (Homo sapiens). The following Table 20 is adapted from our work on vaccine study^49^.

**Table 20.**
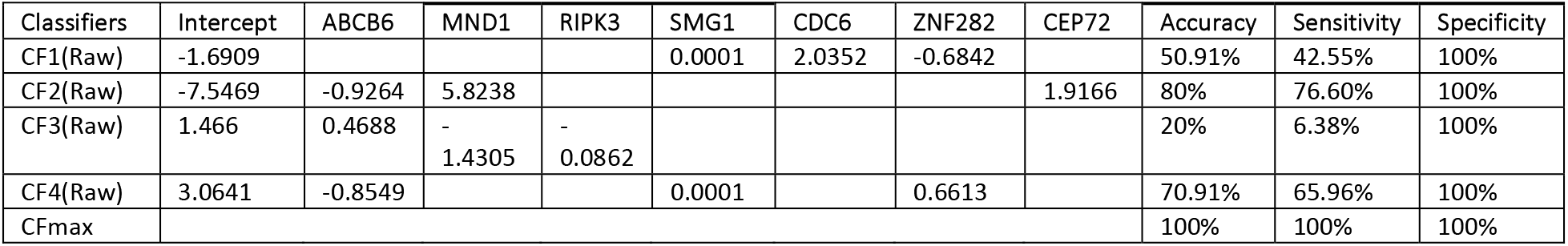
Performance of individual classifiers and combined max-competing classifiers using blood sampled data GSE201530 to classify COVID-19 infected and healthy control into their respective groups. The meaning of CF-i is the same as those in Table 1. Raw stands for raw counts.

It is significant to note that the genes identified from blood samples in our earlier work^17^ again work for various SARS-CoV-2 variants, including Omicron.

## 5 Discussions

The results presented in this paper are the first to directly associate a few critical genes with SARS-CoV-2 with the best performance (relative to other subsets with the same number of genes). Furthermore, the results signify the genomic difference between NP/OP swap PCR samples and whole blood samples (hospitalized patients), identify single-digit critical genes (ATP6V1B2, IFI27, BTN3A1, SERTAD4, EPSTI1), which are a transcriptional response to SARS-CoV-2, interpretable functional effects of gene-gene interactions, gene-variants interactions using explicitly mathematical expressions, introduce graphical tools for medical practitioners to understand the genomic signature patterns of the virus, make suggestions on developing more efficient vaccines and antiviral drugs, and finally identify potential genetic clues to other diseases due to COVID-19 infection.

In Zhang^17^, a conceptual visualization of the gene-gene relationship was created. At the top of the figure, virus variants were placed. With the new findings of this paper, six signature patterns from Tables 3–5 can be used to replace those virus variants, and then a complete dynamic flow can be formed.

As discussed in Introduction, the genes identified in Zhang^17^ are hypothesized to link to the root cause of COVID-19, while the genes identified in this study are the key to treating the symptoms. Therefore, based on the findings in this paper, we make the following hypotheses.

- **Hypothesis 1:** The five genes^17^ ABCB6, KIAA1614, MND1, SMG1, RIPK3, and their functional effects are the key to curing the root cause.
- **Hypothesis 2:** The five genes ATP6V1B2, IFI27, BTN3A1, SERTAD4, EPSTI1, and their functional effects are the key to treating the symptoms.
- **Hypothesis 3:** The genes CDC6 (cell division cycle 6)^17^ and MND1 are protein essentials for the initiation of RNA replication.

Hypothesis 1 is based on the mathematical and biological equivalence between COVID-19 disease, and the functional effects of these five genes proved in Zhang^17^. At the moment, testing Hypothesis 2 is more urgent than testing Hypothesis 1 given variants of SARS-CoV-2 have been emerging, and waves of COVID-19 have been arriving one after another. Furthermore, once Hypothesis 2 is tested and confirmed, scientists can test their counterparts from animals, trace the virus origin, and find the intermediate host species of SARS-CoV-2. As to Hypothesis 3, in Zhang (2021)^17^, a combination of CDC6 and ZNF282 (Zinc Finger Protein 282) can lead to 97.62% accuracy (98% sensitivity, 96.15% specificity), with the following classifier: 1.7615 + 6.8226*CDC6 −1.1556*ZNF282, which suggests the protein encoded by CDC6 is a protein essential for the initiation of RNA replication. In addition, ZNF282 can be a repressor of COVID-19 RNA replication.

As mentioned in Introduction, ATP6V1B2 was found to impair lysosome acidification and cause dominant deafness-onychodystrophy syndrome^25^, while IFI27 was found to discriminate between influenza and bacteria in patients with suspected respiratory infection^24^. There have been new concerns with COVID-19 disease, e.g., SARS-CoV-2 enters the brain^12^, COVID-19 vaccines complicate mammograms^13^, memory loss and ‘brain fog’^14^, SARS-CoV-2 can persist for months after traversing body^26^. Using the findings from this paper, we may hypothesize that ATP6V1B2 can be a leading factor causing COVID-19 to brain function and ENT problems. As to IFI27, given that COVID-19 is a respiratory tract infection, it makes sense to hypothesize IFI27 is the infection’s key. EPSTI1 has been found related to breast cancer and oral squamous cell carcinoma (OSCC) and lung squamous cell carcinoma (LSCC)^35^, which may link COVID-19 to what has been found in mammograms complication^13^. Liang et al. (2021)^36^ suggests that BTN3A1 may function as a tumor suppressor and may serve as a potential prognostic biomarker in NSCLCs and BRCAs. However, all of these findings have not been confirmed. A confirmed Hypothesis 2 may help further explore whether these genes reported in the literature are truly effective, as suggested in the literature.

Finally, with the proven existence of signature patterns associated with SARS-CoV-2 and COVID-19, variants of the disease will continue to emerge if the problems revealed by the existing signatures are not solved. We have witnessed that each time after a peak of the COVID-19 pandemic, the world saw hopes of the end of the pandemic, and the public lowered their guard; as a result, another wave (small or big) appeared. As such, we shouldn’t forget the pain where the gain follows as existence determines recurrence noted by Murphy’s law “Anything that can go wrong will go wrong.”

## Acknowledgments

To be added later

## Supplementary materials

Real data and computer outputs are in a supplementary file available online https://pages.stat.wisc.edu/~zjz/BHDataCode.zip. In addition, a MATLAB^®^ demo code for solving a final dataset example in Equation (4) (*λ*_2_=0) is also available.

## Data Availability

The datasets are publicly available. The data links are stated in Section Data Description.

## Limitation statements

Although we have identified functional effects by gene-gene interactions and gene-subtype (variants) interactions of the five genes, we haven’t identified how gene-gene interacts with each other and their causal directions. We are working in this direction. Finally, our results are in the field of computational biology/medicine, and they are not lab-confirmed.

## References

[1] C. Rowland. Doctors and nurses want more data before championing vaccines to end the pandemic: Health systems are launching bids to assure their medical workers that vaccines will be safe and effective. CNN, pages November 21, 2020 at 6:00 a.m. CST, 2020. URL https://www.washingtonpost.com/business/2020/11/21/vaccines-advocates-nurses-doctorscoronavirus/.

[2] Ewen Callaway. The quest to find genes that drive severe covid. Nature, pages 346–348, 2021. ISSN 595(7867). doi: 10.1038/d41586-021-01827-w.

[3] COVID-19 Host Genetics Initiative. Mapping the human genetic architecture of COVID-19. Nature, pages 1476–4687, 2021. doi: 10.1038/s41586-021-03767-x. URL https://doi.org/10.1038/s41586-021-03767-x.

[4] E. Pairo-Castineira, S. Clohisey, L. Klaric, and et al. Genetic mechanisms of critical illness in COVID-19. Nature, 591:92–98, 2021. URL https://doi.org/10.1038/s41586-020-03065-y.

[5] The RECOVERY Collaborative Group. Dexamethasone in hospitalized patients with covid-19. New England Journal of Medicine, 384(8):693–704, 2021. doi: 10.1056/NEJMoa2021436. URL https://doi.org/10.1056/NEJMoa2021436.

[6] Gillian S. Dite, Nicholas M. Murphy, and Richard Allman. Development and validation of a clinical and genetic model for predicting risk of severe COVID-19. Epidemiology and Infection, 149:e162, 2021.doi: 10.1017/S095026882100145X.

[7] Zhang, Q. et al. Inborn errors of type i ifn immunity in patients with life-threatening COVID-19. Science, 370:eabd4570, 2020. doi: 10.1126/science.abd4570.

[8] Bastard, P. et al. Autoantibodies against type i ifns in patients with life-threatening COVID-19. Science, 370:eabd4585, 2020. doi: 10.1126/science.abd4585.

[9] Povysil, Gundula et al. Failure to replicate the association of rare loss-of-function variants in type IIFN immunity genes with severe COVID-19. medRxiv, 2020. doi: 10.1101/2020.12.18.20248226. URL https://www.medrxiv.org/content/early/2020/12/21/2020.12.18.20248226.

[10] Kosmicki, J. A. et al. Genetic association analysis of SARS-CoV-2 infection in 455,838 UK biobank participants. medRxiv, 2020. doi: 10.1101/2020.10.28.20221804. URL https://www.medrxiv.org/content/early/2020/11/03/2020.10.28.20221804.

[11] Fallerini, Chiara et al. Association of toll-like receptor 7 variants with life-threatening COVID-19 disease in males: findings from a nested case-control study. eLife, 10:e67569, mar 2021. ISSN 2050-084X. doi: 10.7554/eLife.67569.

[12] Elizabeth M. Rhea, Aric F. Logsdon, Kim M. Hansen, and et al. The s1 protein of SARS-CoV-2 crosses the blood{brain barrier in mice. Nature Neuroscience, 24(3):368–378, 2021. URL https://doi.org/10.1038/s41593-020-00771-8.

[13] COVID-19 vaccines complicate mammograms. Cancer Discovery, 11(8):1868–1868, 2021. ISSN 2159-8274. doi: 10.1158/2159-8290.CD-NB2021-0366. URL https://cancerdiscovery.aacrjournals.org/content/11/8/1868.1.

[14] Jacqueline H. Becker, Jenny J. Lin, Molly Doernberg, Kimberly Stone, Allison Navis, Joanne R. Festa, and Juan P. Wisnivesky. Assessment of Cognitive Function in Patients After COVID-19 Infection. JAMA Network Open, 4(10):e2130645–e2130645, 10 2021. ISSN 2574-3805. doi: 10.1001/jamanet-workopen.2021.30645. URL https://doi.org/10.1001/jamanetworkopen.2021.30645.

[15] Zhengjun Zhang. Five critical genes related to seven COVID-19 subtypes: A data science discovery. Journal of Data Science, 19(1):142–150, 2021. https://doi.org/10.6339/21-JDS1005.

[16] Overmyer, Katherine A. et al. Large-scale multi-omic analysis of COVID-19 severity. Cell Systems, page doi.org/10.1016/j.cels.2020.10.003, 2020.

[17] Zhengjun Zhang. The existence of at least three genomic signature patterns and at least seven subtypes of COVID-19 and the end of the disease. Vaccines, 10(5), 761; https://doi.org/10.3390/vaccines100507612022.

[18] Zhengjun Zhang. Lift the veil of breast cancers using 4 or fewer critical genes. Cancer Informatics, 21:1–11, 2022. https://doi.org/10.1177/11769351221076360.

[19] Zhengjun Zhang. Functional effects of four or fewer critical genes linked to lung cancers and new sub-types detected by a new machine learning classifier. Journal of Clinical Trials, 11:S14:100001, 2021. https://www.longdom.org/open-access/functional-effects-of-four-or-fewer-critical-genes-linked-to-lung-cancers-and-new-subtypes-detected-by-a-new-machine-learning-clas-88321.html

[20] Zhengjun Zhang, Yuqing Xu, Xiaoxing Li, Mengke Chen, Xueqin Wang, Ning Zhang, Xiaofei Zhang, Wei Zheng, Heping Zhang, Yongjun Liu. PSMC2 and CXCL8-Modulated Four Critical Gene-Based High-Performance Biomarkers for Colorectal Cancers. Manuscript to be submitted, 2022.

[21] Yongjun Liu, Heping Zhang, Yuqing Xu, Yao-Zhong Liu, Matthew M Yeh, Zhengjun Zhang. The Interaction Effects of GMNN and CXCL12 Built in Five Critical Gene-based High-Performance Biomarkers for Hepatocellular Carcinoma. Manuscript to be submitted, 2022.

[22] E Mick, J Kamm, A.O. Pisco, K Ratnasiri, and et al. Upper airway gene expression reveals suppressed immune responses to SARS-CoV-2 compared with other respiratory viruses. Nat Commun, 11:5854, 2020. URL https://doi.org/10.1038/s41467-020-19587-y.

[23] NAP Lieberman, V Peddu, H Xie, and et al. In vivo antiviral host transcriptional response to SARS-CoV-2 by viral load, sex, and age. PLoS Biol., 18(9):e3000849, 2020.

[24] Benjamin M. Tang, Maryam Shojaei, Grant P. Parnell, and et al. A novel immune biomarker IFI27 discriminates between influenza and bacteria in patients with suspected respiratory infection. European Respiratory Journal, 49(:):1602098, 2017. doi: 10.1183/13993003.02098-2016.

[25] Y Yuan, J Zhang, Q Chang, and et al. De novo mutation in ATP6V1B2 impairs lysosome acidification and causes dominant deafness-onychodystrophy syndrome. Cell Res., 24(11):1370{3, 2014. doi: 10.1038/cr.2014.77.

[26] Daniel Chertow, Sydney Stein, Sabrina Ramelli, and et al. SARS-CoV-2 infection and persistence throughout the human body and brain. Researchsquare, 2021. doi: https://doi.org/10.21203/rs.3.rs1139035/v1

[27] Hao Yang Teng and Zhengjun Zhang. Directly and simultaneously expressing absolute and relative treatment effects in medical data models and applications. Entropy, 23(11), 2021. ISSN 1099-4300. doi: 10.3390/e23111517. URL https://www.mdpi.com/1099-4300/23/11/1517.

[28] J. Aitchison and J. A. Bennett. Polychotomous quantal response by maximum indicant. Biometrika, 57(2):253{262, 08 1970. ISSN 0006-3444. doi: 10.1093/biomet/57.2.253. URL https://doi.org/10.1093/biomet/57.2.253.

[29] Qiurong Cui and Zhengjun Zhang. Max-linear competing factor models. Journal of Business & Economic Statistics, 36(1):62–74, 2018. doi: 10.1080/07350015.2015.1137761.

[30] Qiurong Cui, Yuqing Xu, Zhengjun Zhang, and Vincent Chan. Max-linear regression models with regularization. Journal of Econometrics, 222:579–600, 2021. doi: https://doi.org/10.1016/j.jeconom.2020.07.017.

[31] Daniel McFadden. Econometric Models for Probabilistic Choice among Products. The Journal of Business, 53(3):S13{29, 1980.

[32] Takeshi Amemiya. Advanced Econometrics. Harvard University Press, Cambridge, 1985.

[33] Jing Qin. Discrete Data Models, pages 249{257. Springer Singapore, Singapore, 2017. ISBN 978-981-10-4856-2. doi: 10.1007/978-981-10-4856-2-13. URL https://doi.org/10.1007/978-981-10-4856-2-13.

[34] Zhengjun Zhang. Quotient correlation: a sample based alternative to Pearson’s correlation. The Annals of Statistics, 36(2):1007–1030, 2008.

[35] Mengmeng Fan, Makoto Arai, Akinobu Tawada, and et al. Contrasting functions of the Epithelial stromal interaction 1 gene, in human oral and lung squamous cell cancers. Oncology Reports, 47(1):5, 2022. URL https://doi.org/10.3892/or.2021.8216.

[36] F Liang, C Zhang, Guo H, and et al. Comprehensive analysis of btn3a1 in cancers: mining of omics data and validation in patient samples and cellular models. FEBS Open Bio., 11(9):2586{2599, 2021. doi: 10.1002/2211-5463.13256.

[37] Thair SA, He YD, Hasin-Brumshtein Y, Sakaram S et al. Transcriptomic similarities and differences in host response between SARS-CoV-2 and other viral infections. iScience 2021 Jan 22;24(1):101947. PMID: 33437935

[38] Fong SW, Yeo NK, Chan YH, et al. Whole blood transcriptome analysis reveals SARS-CoV-2 ORF8 as a potential therapeutic and vaccine target.

[39] Ng DL, Granados AC, Santos YA, Servellita V et al. A diagnostic host response biosignature for COVID-19 from RNA profiling of nasal swabs and blood. Sci Adv 2021 Feb;7(6). PMID: 33536218

[40] Vono M, Huttner A, Lemeille S, Martinez-Murillo P et al. Robust innate responses to SARS-CoV-2 in children resolve faster than in adults without compromising adaptive immunity. Cell Rep 2021 Oct 5;37(1):109773. PMID: 34587479

[41] Wang D, Wang D, Huang M, Transcriptomic characteristics and impaired immune function of patients who retest positive for SARS-CoV-2 RNA

[42] Singh NK, Srivastava S, Zaveri L, Bingi TC et al. Host transcriptional response to SARS-CoV-2 infection in COVID-19 patients. Clin Transl Med 2021 Sep;11(9):e534. PMID: 34586723

[43] Masood KI, Yameen M, Ashraf J, Shahid S et al. Upregulated type I interferon responses in asymptomatic COVID-19 infection are associated with improved clinical outcome. Sci Rep 2021 Nov 25;11(1):22958. PMID: 34824360

[44] Galván-Peña S, Leon J, Chowdhary K, Michelson DA et al. Profound Treg perturbations correlate with COVID-19 severity. Proc Natl Acad Sci U S A 2021 Sep 14;118(37). PMID: 34433692

[45] Chen J, Guo M, Identifying Risk Factors for Secondary Infection Post-SARS-CoV-2 Infection.

[46] Knabl L, Lee HK, Wieser M, Mur A et al. BNT162b2 vaccination enhances interferon-JAK-STAT-regulated antiviral programs in COVID-19 patients infected with the SARS-CoV-2 Beta variant. Commun Med (Lond) 2022;2(1). PMID: 35465056

[47] Lee HK, Knabl L, Walter M, Knabl L Sr et al. Prior Vaccination Exceeds Prior Infection in Eliciting Innate and Humoral Immune Responses in Omicron Infected Outpatients. Front Immunol 2022;13:916686. PMID: 35784346

[48] Lee HK, Knabl L, Knabl L Sr, Wieser M et al. Immune transcriptome analysis of COVID-19 patients infected with SARS-CoV-2 variants carrying the E484K escape mutation identifies a distinct gene module. Sci Rep 2022 Feb 18;12(1):2784. PMID: 35181735

[49] Zhengjun Zhang, Genomic Benefits and Potential Harms of COVID-19 Vaccines Indicated from High-Performance Genomic Biomarkers, 2022, manuscript submitted.

